# Reverse Spatiotemporal Hierarchy during Cross-modal Memory Recall and Imagery

**DOI:** 10.1101/2025.11.26.690759

**Authors:** Yu Hu, Jörn Diedrichsen, Yalda Mohsenzadeh

## Abstract

Recalling past events is often accompanied by mental imagery of those experiences. Based on previous research, this process engages memory- and sensory-related brain areas. However, the underlying spatiotemporal dynamics remain poorly investigated. Here, we used naturalistic videos of audiovisual events and recorded fMRI data during the tasks in which human participants recalled visual contents when hearing associated sounds and recalled sounds when watching silent videos, after they had well memorized the video contents. With time-resolved fMRI multivariate pattern analyses, we observed reverse spatiotemporal hierarchy during the visual memory recall and imagery: the neural activity in primary visual cortex was delayed compared with high-order visual areas. A similar pattern was found during auditory memory recall and imagery, where the bottom-up progression from the mid-level planum temporale to the high-level superior temporal gyrus observed during auditory perception was absent. However, the primary auditory area was not involved, suggesting modality differences in the role of primary sensory areas in corresponding memory recall. We also observed the activity of the hippocampus, the parahippocampal cortex, the retrosplenial cortex, and the precuneus and examined their temporal dynamics. Overall, our study provided both spatial and temporal accounts of neural activity during the cross-modal memory recall and imagery.

## Introduction

When we recall past events, we often create mental imagery to re-experience aspects of the original events. Many people can recreate the visual experience from their memory in their ”mind’s eye”, although some populations cannot^1^. For example, recalling a vacation might bring up visualizations of the visited place. However, memories often include more than just visual elements, and other sensory experiences can also be recreated. For instance, remembering a concert might involve hearing parts of the music or feeling the vibration of the bass.

This process first involves memory recollection, which recruits a network of brain regions to retrieve past information^2–4^. Previous work shows that the hippocampus and the parahippocampal cortex (PHC) in the medial temporal lobe are critical for episodic memory retrieval^5–8^, which is dissociated from familiarity-based recognition mediated mainly by the perirhinal cortex (PRC)^6,9^. Moreover, areas in the default mode network^10,11^, such as the retrosplenial cortex (RSC), the precuneus (PCu), and the ventromedial prefrontal cortex (vmPFC), also play a role in the recall of the past^12–15^.

This process also involves mental imagery to reconstruct past experiences. Many studies show that mental imagery engages sensory brain areas as original sensory perceptions. Visual mental imagery involves the visual cortex to create the internal visual experience^16–19^. The high-order visual areas exhibit a larger overlap between visual perception and imagery^20,21^, in which the left fusiform region is found to be consistently activated across previous studies on visual imagery^22,23^. Some studies reported that the early visual cortex is not significantly activated above baseline^16,23–25^, but studies using multivariate approaches^26–29^ or low-level visual stimuli^30–33^ identified its engagement. Its involvement also depends on the experimental instructions^34^ and imagery vividness^27,35^. Similarly, auditory imagery, although less studied, has also been found to activate the auditory cortex^16,36,37^.

However, little is known about the temporal hierarchy in these brain areas during memory retrieval and imagery. Although previous studies show that the level of information flow is reversed during memory reconstruction or mental imagery^38–41^, we still lack direct evidence characterizing the time course of neural activity in specific brain regions during this process. To this end, we applied time-resolved fMRI multivariate decoding analysis^42^ and aimed to assess whether reverse temporal hierarchy can be observed at the temporal resolution of the fMRI signals in one second. To investigate the memory recall and imagery process regardless of sensory modalities, we used naturalistic audiovisual stimuli and examined neural activity during both visual and auditory memory recall and imagery triggered by associated cross-modal information.

## Results

We curated 80 one-second videos that capture common daily audiovisual events of animals, objects, people, and scenes. We first recorded fMRI data when participants (n=16) passively viewed the videos with accompanying sounds (**Figure 1**). Next, we asked participants to memorize the video contents and repeatedly practice recalling visual contents when hearing only the sounds and recalling sounds when watching silent videos. During each practice, they were asked to create mental imagery as vividly as possible and rate the vividness after each practice. We ended the behavior session only when the accumulated vividness score reached 80% out of a five-point scale. After that, we recorded fMRI data when participants alternatively performed the audio-induced visual memory recall and video-induced auditory memory recall tasks (see details in Methods).

**Figure 1:**
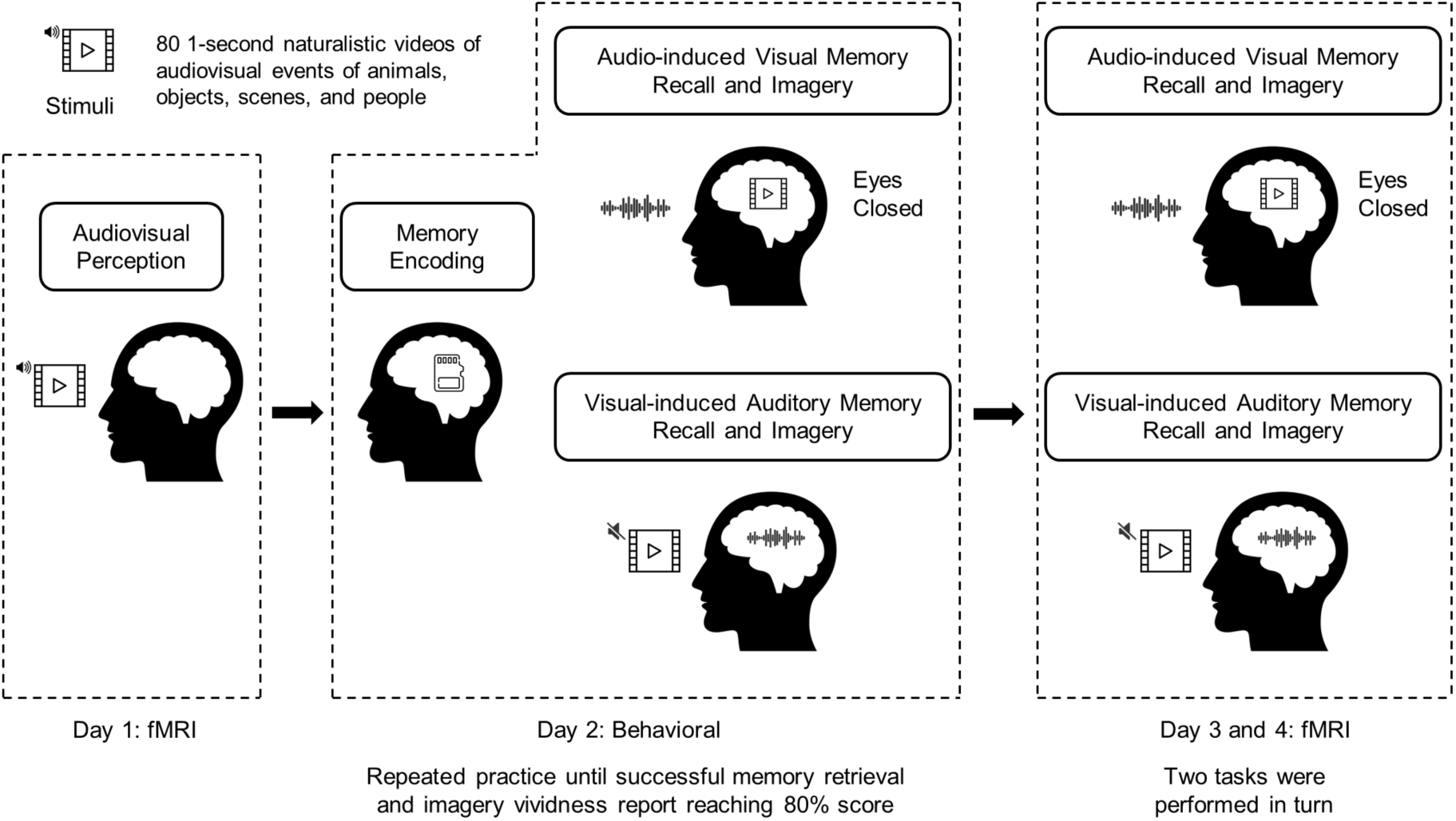
Experimental design. The stimuli used are 80 1-second naturalistic audiovisual video clips that capture common daily audiovisual events of animals, objects, people, and scenes. The whole experiment was conducted on four different days. On day 1, participants (n=16) were scanned with fMRI when they passively viewed videos with orthogonal oddball detection to maintain their attention. On day 2, participants memorized the video contents and practiced recalling visuals when hearing only sounds and recalling sounds when watching silent videos. They were also instructed to create mental imagery as vividly as possible. After each practice, they were asked to report the vividness of imagery. They repeated this process until they could successfully achieve memory recall and the accumulated 5-point vividness score reached 80%. On days 3 and 4, they performed the same memory recall and imagery tasks with the fMRI data collected. To collect enough data trials, the tasks were completed within two days in which the two tasks were performed alternately.

We investigated neural responses during these tasks in brain areas associated with visual and auditory processing as well as memory retrieval. We first assessed the univariate level of activations in contrast to the resting baseline in each region of interest (ROI) using conventional GLM modeling. Next, we applied time-resolved multivariate decoding analyses to examine how the brain activity unfolded over time. Specifically, we followed the spatiotemporal estimation method proposed by Turner et al.^42^ to estimate beta weights at each repetition time (TR=1 second) of fMRI signals. We then performed cross-validation to evaluate the decoding accuracy to classify the stimulus category at each time point using the linear support vector machine (SVM). This approach allowed us to examine the temporal evolution of class-discriminating information in certain brain ROIs.

When testing whether the decoding accuracy were significantly above chance at each time point, we first normalized the accuracy values by the mean of the maximum peak and then performed one-sided 10,000 sign-permutation test to identify the time point at which decoding accuracy exceeded chance (p<0.05, FDR corrected). To quantify the temporal differences between ROIs and address the effect of signal strength on determining the onset of significant decoding^43^, we applied cross-correlation analyses. We first linearly interpolated the values by 100 ms and then identified the temporal shift that yielded the maximum cosine similarity between the curves. For each comparison, we used the averaged decoding accuracy curve of one brain region as the reference and estimated the confidence interval of temporal lags by applying 10,000 bootstrap resampling on the accuracy values of the other area. This procedure was repeated with the two regions switched, and a temporal lag was considered significant only if both ways of comparison yielded significant results. Consequently, we have measures based on the significant values and on the shape of the decoding curves, providing a complementary characterization of the temporal evolution of neural information.

### Reverse hierarchy in visual areas during visual memory recall and imagery

To examine visual regions at different processing levels, we selected primary visual cortex (V1) as a low-level, V4 as a middle-level, the lateral occiptal cortex (LOC) and the fusiform gyrus (FG) as high-level regions (**Figure 2a**). The LOC is commonly assessed as the high-level visual area to represent the object category^44^ and the FG is selected because a meta-analysis shows that it is consistently activated during visual imagery as the fusiform imagery node^23^. We found that during the audiovisual perception task and visual-induced auditory memory recall task, all visual areas examined were significantly activated above baseline (**Figure 2b**). The decoding accuracy for the two tasks rose above chance after 3 seconds for V1, V4 and, LOC, but after 4 seconds for FG (**Figure 2c**). For audiovisual perception task, the estimated temporal lag between V1 and LOC was 0.3 s (95% Confidence interval 0.2-0.4 s), between V1 and FG was 0.6 s (0.4-0.7 s), between V4 and LOC was 0.2 s (0.1-0.4 s), between V4 and FG was 0.5 s (0.3-0.6 s), and between LOC and FG was 0.2 s (0.1-0.4 s) (**Figure 2d**). Only the lag between V1 and V4 was not significantly different than zero. A similar pattern was found for visual-induced auditory memory recall task, where the estimated temporal lag between V1 and LOC was 0.2 s (0.1-0.3 s), between V1 and FG was 0.7 s (0.5-0.9 s), between V4 and LOC was 0.3 s (0.2-0.4 s), between V4 and FG was 0.8 s (0.6-1.0 s), and between LOC and FG was 0.5 s (0.3-0.7 s). For the task of audio-induced visual memory recall and imagery, we did not observe any significant univariate activations in V1, V4 and LOC, except for FG. This is consistent with the meta-analysis showing that only the fusiform area was activated during visual imagery^23^. However, categorical information can be decoded in V1, LOC, and FG, suggesting that the low-level visual area V1 was involved during visual memory recall and imagery even with the low magnitude of activation. However, the middle-level visual area V4 was not involved because no categorical information was decoded from the activity patterns in this region. Prominently, we found that the onset of significant decoding accuracy in V1 was one second delayed compared with high-level visual areas (**Figure 2c**). The estimated temporal shift between V1 and LOC was -0.7 s (-1.0 to -0.3 s), and between V1 and FG was -1.1 s (-2.8 to -0.5 s). No significant difference was observed between LOC and FG (**Figure 2d**). This demonstrate that the V1 was involved later in time after the activity in high-level visual areas.

**Figure 2:**
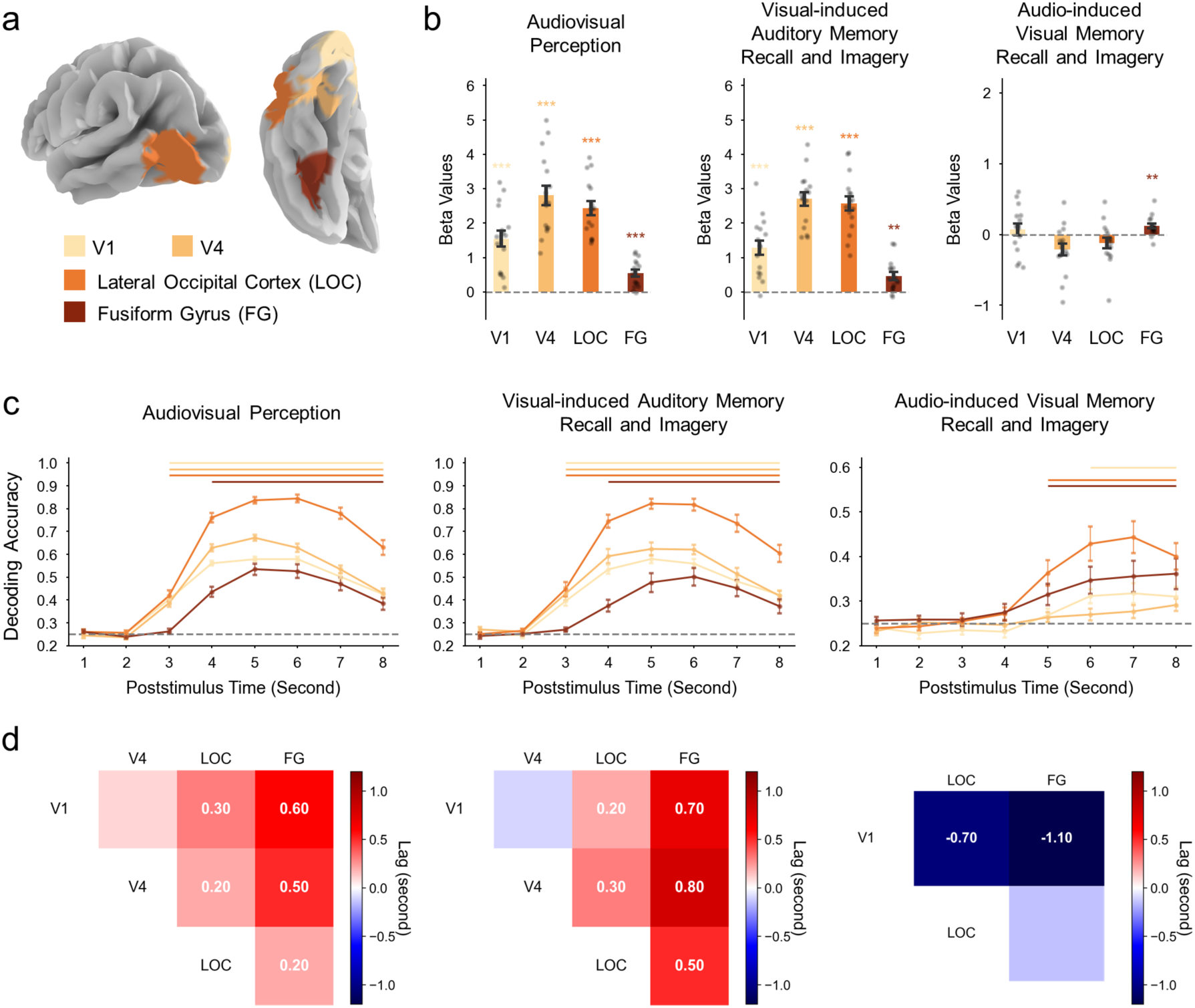
Univariate responses and time-resolved multivariate decoding in visual areas. **(a)** Visualization of visual ROIs. **(b)** Univariate beta responses of visual ROIs for three different tasks. For each task, the beta weights were estimated across all stimuli and averaged within each ROI. The height of the bar denotes the mean beta values averaged across subjects (n=16) for an ROI. The error bars represent the standard error of the mean. The grey dots depict individual values for each subject. The asterisks above denote significant differences above zero based on one-sample t-tests (*p<0.05, **p<0.01, ***p<0.001). The color corresponds to different ROIs. **(c)** Time-resolved multivariate categorical decoding of visual ROIs. For each task, the beta weights were estimated for each stimulus and at each TR time (1 second) using spatiotemporal estimation method^42^. Next, at each time point, the beta values for 80 stimuli within an ROI were used to classify the stimuli category through the linear support vector machine (SVM) and the cross-validation method. This yielded a time course of decoding accuracy for each subject and each ROI. The gray dashed line denotes the value of chance accuracy. The time course of dot values indicates the mean decoding accuracy averaged across subjects at each time point, and the error bars represent the standard error of the mean. The color corresponds to different ROIs. The lines above denote time points of significant above-chance accuracy. To determine the significant time points, the accuracy values were first normalized by the mean of the maximum peak and one-sided 10,000 sign-permutation test was performed at each time point (p<0.05, FDR corrected). **(d)** Estimated temporal lags of decoding curves between pairs of visual ROIs. If two ROIs both showed significant decoding accuracy, we computed their temporal shifts by the cross-correlation method. We linearly interpolated the time course of accuracy values to 100 ms resolution and identified the temporal shift that yielded the maximum cosine similarity between the two curves. For each comparison, the subject-averaged decoding accuracy time course from one brain region was used as the reference. The accuracy values from another region was then resampled 10,000 times across subjects to estimate the median and 95% confidence interval (CI) of the temporal lag between the two regions. This procedure was repeated with the two regions switched, and a temporal lag was considered significant only if both ways of comparison yielded significant results. In the matrix, the color denotes the medium of temporal lag between two regions. A positive lag indicates that the region in the column was delayed relative to the region in the row, whereas a negative lag indicates the opposite. Numerical values are displayed only when the 95% CI did not include zero, indicating a statistically significant temporal difference.

### Absence of bottom-up hierarchy in auditory areas during auditory memory recall and imagery

Similarly, we examined different levels of auditory regions for the three tasks. We selected the Heschl’s gyrus (HG) as a low-level, the planum temporale (PT) as a middle-level, and the posterior superior temporal gyrus (pSTG) as a high-level auditory area (**Figure 3a**). During the audiovisual perception and audio-induced visual memory recall task, we observed significant activations in these auditory ROIs (**Figure 3b**). The decoding accuracy in all three ROIs became significant at 3 s for the audiovisual task and at 4 s for the audio-induced visual memory recall task(**Figure 3c**). The difference in activation magnitude and decoding onset between the two tasks may reflect the effect of multisensory modulations^45–47^. For the perception task, the temporal lag between HG and PT was 0.4 s (0.2-0.5s), between HG and pSTG was 0.5 s (0.4- 0.6 s), and between PT and STG was 0.2 (0.1-0.3 s) (**Figure 3d**). For the audio-induced visual memory recall task, the temporal lag between HG and PT was 0.6 s (0.4-0.7s), between HG and pSTG was 0.9 s (0.6-1.1 s), and between PT and STG was 0.2 (0.1-0.3 s). All lag values were significantly different than zero, suggesting the bottom-up hierarchical progression. During auditory memory recall and imagery, we found no involvement of HG because no significant positive univariate activation or categorical decoding was observed. Instead, we observed significant decoding accuracy in PT and pSTG, despite no significant univariate activations above baseline. The significant decoding onset of PT was one second delayed compared with that of pSTG (**Figure 3c**). However, the temporal shift between the two areas was not statistically significant (**Figure 3d**). Although this lack of a significant shift does not provide direct evidence for a reverse hierarchy, the contrast with the other two tasks suggests that the bottom-up temporal order observed during perception was absent during auditory memory recall and imagery.

**Figure 3:**
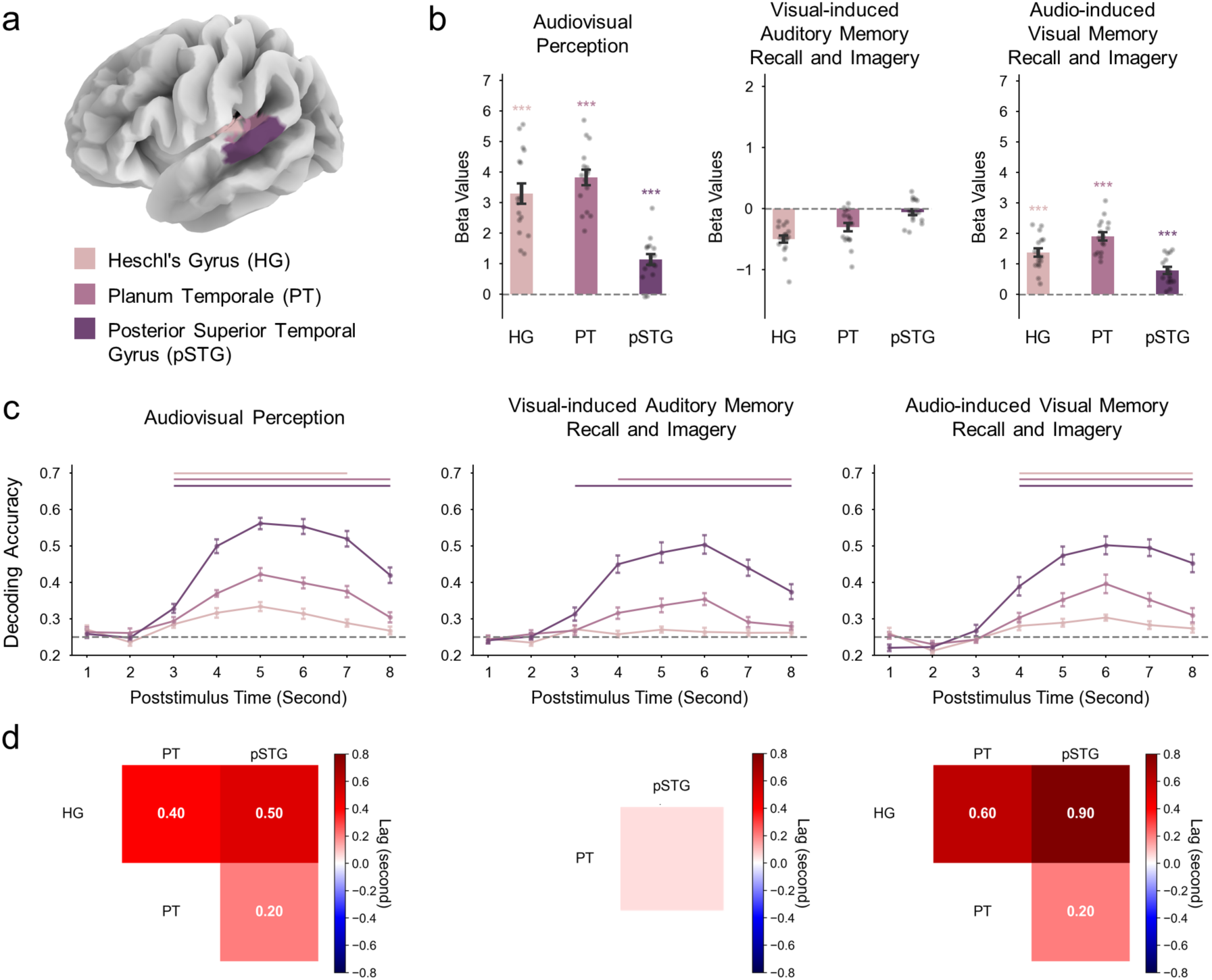
Univariate responses and time-resolved multivariate decoding in auditory areas. **(a)** Visualization of auditory ROIs. **(b)** Univariate beta responses of auditory ROIs for three different tasks. For each task, the beta weights were estimated across all stimuli and averaged within each ROI. The height of the bar denotes the mean beta values averaged across subjects (n=16) for an ROI. The error bars represent the standard error of the mean. The grey dots depict individual values for each subject. The asterisks above denote significant differences above zero based on one-sample t-tests (*p<0.05, **p<0.01, ***p<0.001). The color corresponds to different ROIs. **(c)** Time-resolved multivariate categorical decoding of auditory ROIs. The gray dashed line denotes the value of chance accuracy. The time course of dot values indicates the mean decoding accuracy averaged across subjects at each time point, and the error bars represent the standard error of the mean. The color corresponds to different ROIs. The lines above denote time points of significant above-chance accuracy. To determine the significant time points, the accuracy values were first normalized by the mean of the maximum peak and one-sided 10,000 sign-permutation test was performed at each time point (p<0.05, FDR corrected). **(d)** Estimated temporal lags of decoding curves between pairs of auditory ROIs. If two ROIs both showed significant decoding accuracy, we computed their temporal shifts by the cross-correlation method. In the matrix, the color denotes the medium of temporal lag between two regions estimated by bootstrapping. A positive lag indicates that the region in the column was delayed relative to the region in the row, whereas a negative lag indicates the opposite. Numerical values are displayed only when the 95% confidence interval does not include zero, indicating a statistically significant temporal difference.

### Temporal dynamics in the medial temporal and parietal cortex

We selected the hippocampus, the parahippocampal cortex (PHC), the retrosplenial cortex (RSC), and the precuneus (PCu) as ROIs associated with memory retrieval (**Figure 4a**). Because the resting state is not a proper baseline to examine the univariate activations during memory tasks in the medial temporal lobe^48^ and we did not have a valid task serving as the contrast, we did not interpret the univariate activations in these areas. Using multivariate approaches, we found that categorical information can be successfully decoded in all examined areas for three different tasks (**Figure 4b**). Although we did not ask participants to purposefully memorize the videos during the perception task, these ROIs were also involved, possibly reflecting the spontaneous memory encoding process. Unlike the results in sensory areas, the significant decoding onset and the temporal shift based on the decoding curves among these four ROIs did not yield consistent results. Regarding the onset time of significant decoding in these ROIs during two memory recall tasks, we found that the temporal patterns were consistent across two sensory modalities. The decoding accuracy in PHC and Pcu became significant first, at the same second with the high-order sensory areas (LOC and FG for visual recall, pSTG for auditory recall). The significant onset in the hippocampus and RSC was delayed for one second, at the same timing with the lower-level sensory areas (V1 for visual recall, PT for auditory recall). In contrast, the temporal order of the significant onset in these ROIs during audiovisual perception, and its relative timing compared to sensory regions, were distinct from those during memory recall. The significant onset of PHC was in the same second with all sensory areas, while the significant onset of the hippocampus and Pcu was delayed for 1 second, and the significant onset of RSC was delayed for 2 seconds. However, when examining the temporal lag between the decoding curves of these ROIs, we only found that RSC was significantly delayed compared to all other ROIs for the audiovisual perception task and the visual-induced auditory memory recall task (**Figure 4c**). For the perception task, the temporal lag between PHC and RSC was 0.9 s (0.2-1.5s), between hippocampus and RSC was 1.3 s (0.5-2.3 s), and between Pcu and RSC was 0.7 (0.1-1.3 s). For the visual-induced auditory memory recall task, the temporal lag between PHC and RSC was 0.7 s (0.2-1.3 s), between hippocampus and RSC was 0.8 s (0.3-1.5 s), and between Pcu and RSC was 0.8 (0.3-1.4 s). In contrast, for the audio-induced visual memory recall task, no significant temporal shift was observed for each pair of the four ROIs.

**Figure 4:**
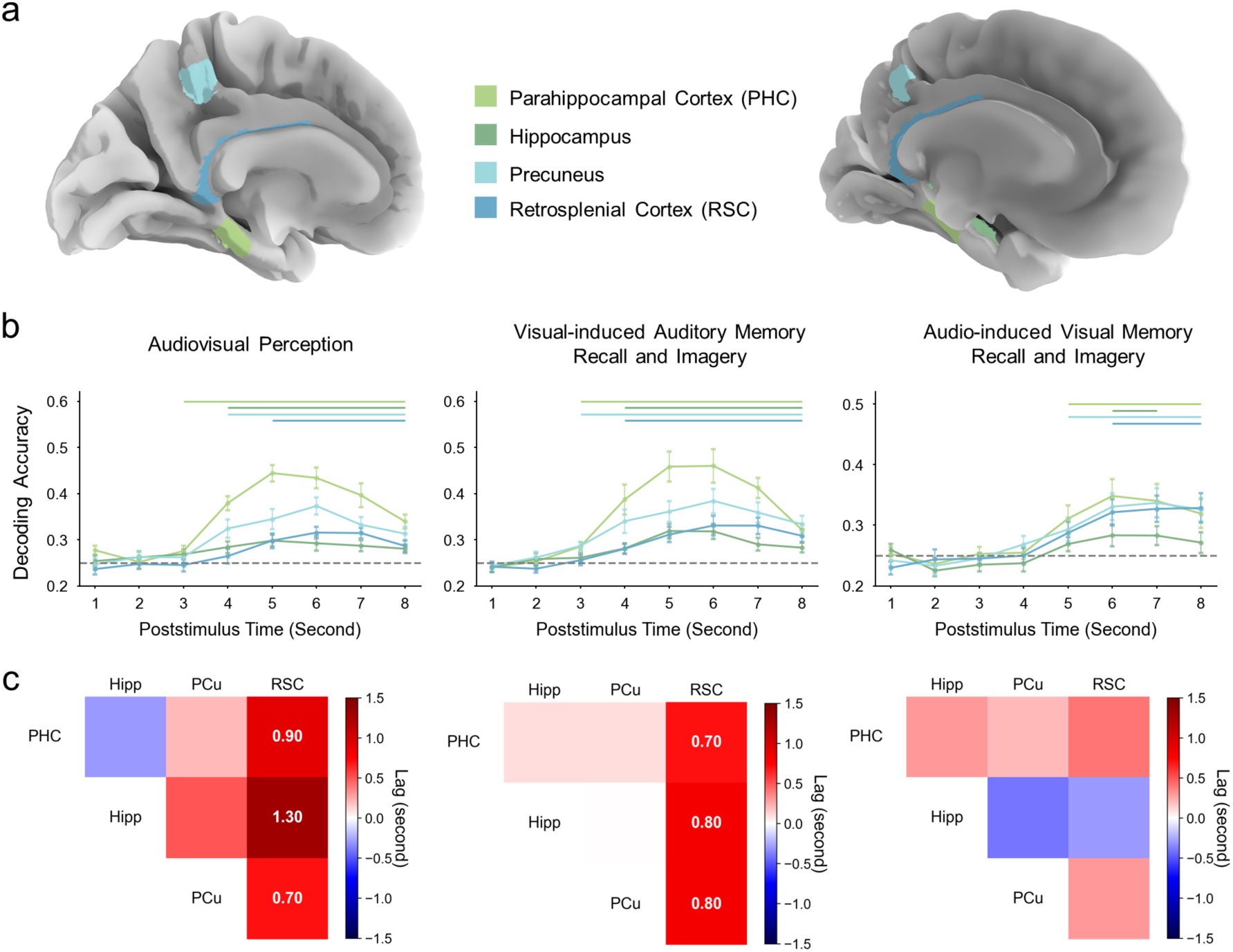
Time-resolved multivariate decoding in the medial temporal and parietal areas. **(a)** Visualization of ROIs associated with memory retrieval in the medial temporal and parietal areas. **(b)** Time-resolved multivariate categorical decoding in these ROIs. For each task, the beta weights were estimated for each stimulus and at each TR time (1 second) using spatiotemporal estimation method^42^. Next, at each time point, the beta values for 80 stimuli within an ROI were used to classify the stimuli category through the linear support vector machine (SVM) and the cross-validation method. This yielded a time course of decoding accuracy for each subject and each ROI. The gray dashed line denotes the value of chance accuracy. The time course of dot values indicates the mean decoding accuracy averaged across subjects at each time point, and the error bars represent the standard error of the mean. The color corresponds to different ROIs. The lines above denote time points of significant above-chance accuracy. To determine the significant time points, the accuracy values were first normalized by the mean of the maximum peak and one-sided 10,000 sign-permutation test was performed at each time point (p<0.05, FDR corrected). **(c)** Estimated temporal lags of decoding curves between pairs of ROIs associated with memory retrieval. If two ROIs both showed significant decoding accuracy, we computed their temporal shifts by the cross-correlation method. In the matrix, the color denotes the medium of temporal lag between two regions estimated by bootstrapping. A positive lag indicates that the region in the column was delayed relative to the region in the row, whereas a negative lag indicates the opposite. Numerical values are displayed only when the 95% confidence interval does not include zero, indicating a statistically significant temporal difference.

### Temporal differences between visual, auditory, and memory-related areas

We next examined the temporal differences between decoding curves across visual, auditory, and memory-related ROIs (**Figure 5**). For the audiovisual perception task, no significant temporal shift was observed between low- and middle-level visual areas (V1 and V4 respectively) and low-level and middle-level auditory areas (HG and PT respectively). The high-level visual areas LOC and FG was significantly delayed relative to HG, and FG was also delayed relative to PT. The high-level auditory area pSTG was significantly delayed relative to both V1 and V4. Among the memory-associated regions, RSC was significantly delayed compared to all visual and auditory areas, while PHC and PCu were significantly delayed relative to auditory area HG and visual areas V1 and V4.

**Figure 5:**
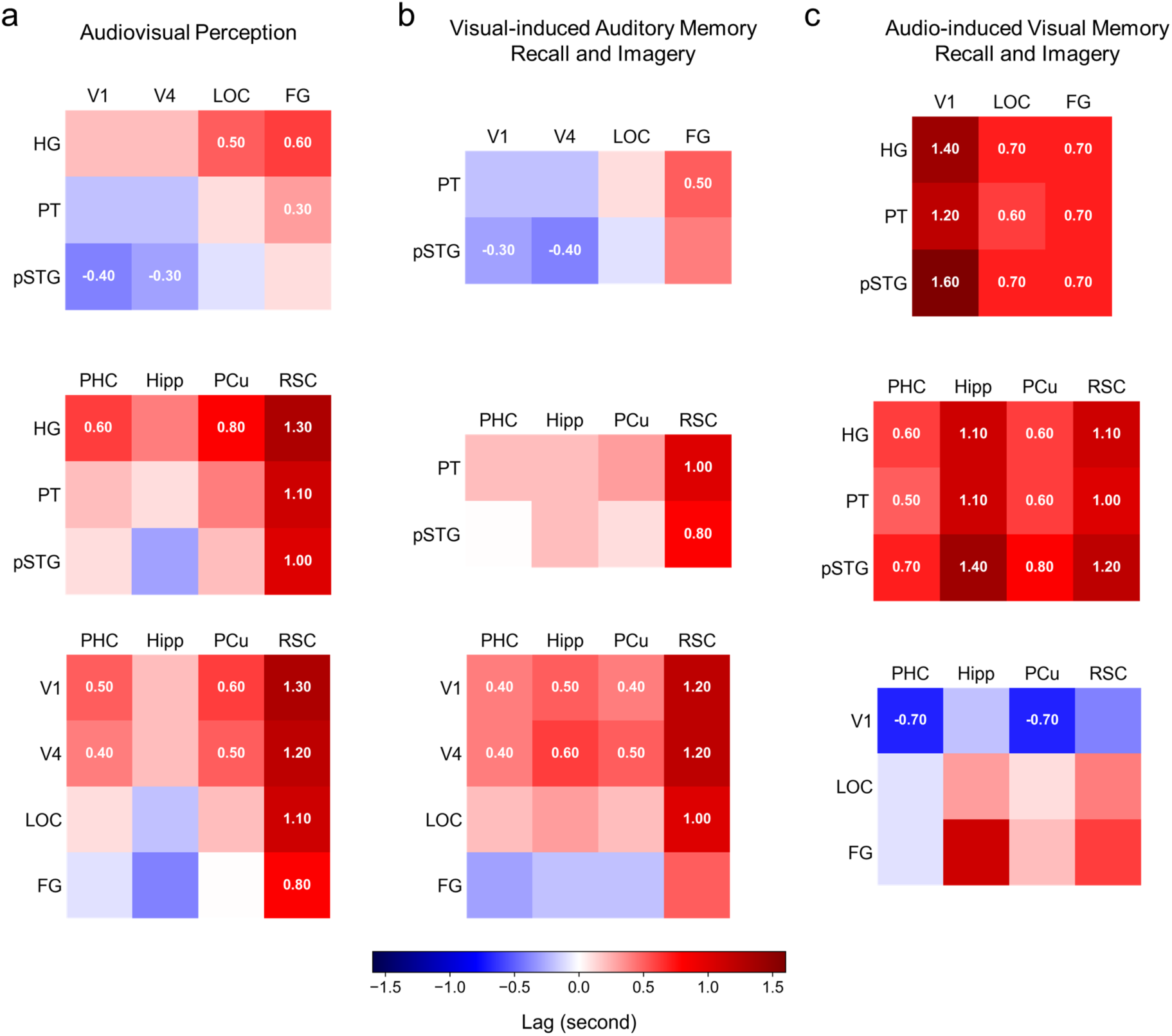
Temporal lags of decoding curves among visual, auditory, and memory ROIs. For each task, if examined ROIs showed significant decoding accuracy, the temporal shifts were estimated among visual, auditory, and memory ROIs. In the matrix, the color denotes the medium of temporal lag between two regions estimated by bootstrapping. A positive lag indicates that the region in the column was delayed relative to the region in the row, whereas a negative lag indicates the opposite. Numerical values are displayed only when the 95% confidence interval does not include zero, indicating a statistically significant temporal difference.

For the visual-induced auditory memory recall task, the pattern of temporal shifts between visual and auditory areas did not differ significantly from that observed during perception. RSC was significantly delayed compared to all visual and auditory areas except for FG. PHC, PCu and hippocampus were significantly delayed relative to visual areas V1 and V4 but not to auditory areas.

For the audio-induced visual memory recall task, all visual regions were significantly delayed compared to auditory regions, with V1 showing the largest temporal lag. All memory-related areas were significantly delayed relative to all auditory areas, but PHC and PCu significantly preceded visual areas V1.

## Discussion

Memory recall is often accompanied by mental imagery. This process engages the medial temporal and parietal areas to retrieve past information^3,14^ and the sensory areas to reconstruct past experiences^16,21^. However, the temporal dynamics in these regions remain largely unexplored. In this study, we employed time-resolved decoding analysis on fMRI data to characterize the temporal hierarchy of neural activity in specific brain areas at a resolution of one second. To assess memory recall across different sensory modalities, we investigated both visual and auditory memory recall and imagery using naturalistic audiovisual stimuli.

Our study first examined the univariate activations in the sensory areas during memory recall and imagery processes. Previous studies assessing univariate activations in early visual areas during visual imagery reported mixed results, potentially due to variations in task instructions^34^, stimulus types, and the imagery vividness^27,35^. Memory type may also be a contributing factor, as working memory tends to preserve more detailed visual information than long-term memory^49^, leading to more vivid imagery and stronger activations. One previous fMRI study^50^ examined the brain responses during both visual and auditory memory recall and imagery using pairs of images and sound. Similarly to their results, we also did not found significant activations in V1 but in fusiform areas. Furthermore, our results align with a meta-analysis^23^ and recent empirical work^51^, highlighting the fusiform region as a reliable site of activation during visual imagery. However, in contrast to^50^, which reported significant activation in secondary auditory areas during auditory memory recall and imagery, we did not observe such activations. This discrepancy may be due to differences in experimental design. Importantly, univariate signal strength alone does not fully capture the involvement of a brain region and its multivariate activity patterns may still encode relevant information despite low activation magnitudes^19^.

Our results revealed opposite temporal dynamics in visual regions during visual memory reconstruction compared to visual perceptual processing. During the tasks of audiovisual perception and visual-induced auditory memory recall, where visual perception is directly engaged, we found that high-level visual areas LOC and FG were involved later than V1. This temporal sequence aligns with the well-established hierarchical progression of visual processing, where information flows from early to higher-order visual areas^52–54^. In contrast, during visual memory recall and imagery, we observed the reverse pattern that LOC and FG were engaged earlier compared to V1, consistent with the hypothesis that visual imagery recruits a top-down hierarchy from high-order to early visual areas^21,41^.

One previous study^40^ showed that the semantic features are processed earlier than perceptual features during object memory reconstruction, which was the opposite order of object perception. Other visual imagery studies^39,41,55^ found that the information represented in EEG signals or the direction of connectivity during perception was reversed during visual imagery. Although these findings suggest the top-down processing in sensory areas during memory reconstruction, no direct evidence is linked to specific brain regions. Our results instead provided the direct evidence characterizing the temporal hierarchy in specific brain areas, revealing the top-down activity during visual memory recall and imagery.

For auditory areas, we found that during the tasks of audiovisual perception and audio-induced visual memory recall, where auditory perception is engaged, the HG was involved first, followed by PT and then pSTG. This temporal progression is consistent with previous findings on the hierarchical organization and temporal dynamics of auditory processing^56–58^. In contrast, during auditory memory recall and imagery, no significant temporal differences were observed between the PT and pSTG. Although we did not find clear evidence for a top-down hierarchy in these regions, the absence of a bottom-up progression suggests that auditory areas follow a distinct temporal pattern during auditory memory reconstruction compared to auditory perception.

Our findings also suggest a modality difference in the involvement of primary sensory areas. We identified the involvement of V1 during visual memory recall but not HG during auditory memory recall. This aligns with prior studies on auditory imagery reporting no activations in the primary auditory cortex^36,37,59,60^. One possible explanation for this discrepancy between modalities is that visual imagery benefits from a retinotopic representation in V1, whereas auditory imagery may rely on more abstract, non-tonotopic representations that do not necessitate A1 involvement. Alternatively, methodological factors such as task design or the sensitivity of our analysis approach may have influenced the observed neural activation patterns. Future research could further investigate these modality-specific differences by employing high-resolution fMRI and advanced analyses to determine the precise neural mechanisms underlying sensory-specific memory recall.

We also observed that the neural processing of visual recall triggered by auditory information unfolded more slowly than auditory recall triggered by visual input. During auditory recall, the high-level auditory area pSTG did not show significant temporal differences compared to the high-level visual areas. However, the high-level visual regions were significantly delayed relative to auditory areas. This temporal asymmetry likely reflects modality-specific retrieval demands. Recalling visual content requires accessing detailed spatial layouts, object features, and scene context, which entails greater cognitive effort and longer buildup time. In contrast, auditory representations may be accessed more rapidly, possibly because sound patterns are inherently sequential and less spatially complex, allowing for faster reactivation of stored information.

We identified the involvement of areas in medial temporal and parietal cortices for three tasks, but we only found RSC showed a significantly delayed temporal lag relative to other memory ROIs during the audiovisual perception and visual-induced auditory memory recall task, but not during the audio-induced visual memory recall task. The delay suggests that RSC may be recruited later in the processing stream when the task involves integrating or reconstructing complex multimodal information. For the task of audiovisual perception and visual-induced auditory memory recall task, we did not observe that the memory-related areas were engaged at different timing compared to high-level visual and auditory areas, except for the RSC. However, during the task of audio-induced visual memory recall, the memory-related areas were involved later compared to auditory areas, at similar timing with high-level visual areas, followed by the V1. Such distinction again suggests that visual recall and reconstruction took a longer time to build up compared to auditory recall.

Still, a primary limitation of this study lies in the temporal resolution of fMRI. Although our analyses leveraged time-resolved multivariate decoding to track the evolution of neural representations, the intrinsic temporal constraints of the hemodynamic response limit the precision with which we can capture rapid neural dynamics. As a result, the temporal lags we observed should be interpreted as coarse approximations rather than precise estimates of the underlying neural timing. Future studies combining fMRI with high-temporal-resolution techniques, such as magnetoencephalography (MEG) or electroencephalogram (EEG), could help disentangle these rapid processes and provide a more complete characterization of the temporal hierarchy of perception and memory reconstruction.

## Methods

### Participants

In total, we recruited 21 participants. When recruiting participants, we asked them to complete the Vividness of Visual Imagery Questionnaire (VVIQ)^61^to assess their self-reported ability to create visual imagery. If the average score of the VVIQ was below 3, we did not recruit the participant. However, we did not inform them that there was a selection process so that they would not intend to cheat to be recruited. Among 21 subjects, 2 withdrew for personal reasons, 1 withdrew because of nausea after the MRI scan, 1 withdrew because of large head movement caused by a cough during the scan, and 1 withdrew because of falling asleep. Consequently, we ended up with 16 subjects (11 females; mean age = 23.25, SD = 3.68). The mean value of VVIQ is 3.68. All participants were right-handed with normal or corrected-to-normal vision and normal hearing. They reported no history of neurological or psychiatric disorders and provided written consent. The study was approved by the Non-Medical Research Ethics Board at Western University and all participants were compensated for their time. All ethical regulations relevant to human research participants were followed.

### Stimuli

We curated 20 naturalistic audiovisual video clips for each category of daily common objects, animals, people, and scenes (see Supplementary Table). Each video clip includes representative visuals and sounds of a certain event. When curating the stimuli, we first found high-quality videos online for a specific event and cropped each one into one second with a frame rate of 25 per second and a frame size of 1920 x 1080. If the video contained no audio or low-quality sounds, we found high-quality audio from other sources and made sure the video and audio were matched and synchronized. We also ensured that the audio contained only clear sound for the specific event without any unrelated sounds such as background noise and the audio alone can be recognized. The audios were normalized root mean squared and resampled to 48 kHz.

### Procedure

The experiment consisted of four sessions completed on four different days but not necessarily consecutive days. We required participants to schedule the experiment dates within two weeks based on their own availability and the availability of MRI scanners.

In the first session, participants were scanned with fMRI when they passively viewed the videos with accompanying sounds. An orthogonal oddball detection task was used to maintain their attention. This session contained eleven runs. In each run, 80 stimuli were played randomly with 10 noise videos in between. The participants were asked to press the button when detecting the noise video. In each trial, the stimuli were played for one second followed by a jittered interstimuli interval in the range of 2-6 seconds.

In the second session, participants were asked to memorize the videos and practice recalling visual contents when only hearing the sounds and recalling sounds when watching silent videos. They were also asked to create imagery as vivid as possible when recalling either the visual or auditory contents. This behavior session included two parts. In the first part, they were allowed to freely go over each video and practice as many times as possible. The second part was composed of multiple runs. In each run, either sounds or silent videos of the stimuli were played randomly, and participants needed to recall the cross-modal contents. After each trial, they were asked to rate the vividness of the mental imagery and were allowed to replay the video to strengthen their memory. They needed to complete at least two runs for each of the two tasks. If the accumulated score of vividness for each video did not reach 80%, they needed to practice more runs until the threshold was reached.

In the next two sessions, participants were scanned with fMRI when they performed the well-practiced sound-induced visual memory recall task and the visual-induced auditory memory recall task. In each session, the two tasks were conducted alternately with six runs for each task. In each run of the sound-induced visual memory recall task, the 80 sounds were played randomly and participants were asked to close their eyes to perform this task. In each trial, after the sound was played for one second, participants were given two seconds to recall the visual and create visual imagery. After that, a beep sound was played and participants were instructed to press the button, stop visual imagery, and simply wait until the next stimulus during a jittered interstimulus interval in the range of 2-6 seconds. The purpose of the beep sound was to restrict the window to perform mental imagery and leave enough resting time between consecutive stimuli, which is essential for the rapid event-related design. Another purpose of the beep sound was to increase the vigilance of the participants and monitor their wakefulness. For the visual-induced auditory memory recall task, the experimental design was similar to the audiovisual perception task in the first session, except for that no sounds were presented. The oddball detection task was also used to maintain their vigilance. We did not leave additional time for auditory imagery as most participants reported that the auditory imagery was simultaneous with the videos.

All experiments were presented through MATLAB Psychtoolbox (http://psychtoolbox. org/). Each video was presented at the center of the screen (view angle *≈* 5°) on a grey background with a black fixation cross. The audios were presented through earphones at a comfortable volume set individually for each participant.

### fMRI data acquisition

Functional MRI data were acquired using a Siemens 3T Magnetom Prisma Fit MRI scanner at the Centre for Functional and Metabolic Mapping at Western University. T2*-weighted, single-shot, gradient-echo echo-planar imaging (GE-EPI) sequence was used with the following parameters: TR = 1000 ms, TE = 30 ms, flip angle = 48°, FOV = 210 mm, multiband factor =5, slice thickness = 2.5 mm, voxel size = 2.5 × 2.5 × 2.5 mm, and 60 axial slices covering the whole brain. High-resolution anatomical images were also acquired using a T1-weighted MPRAGE sequence with the following parameters: TR = 2300 ms, TE = 2.98 ms, flip angle = 9°, FOV = 256 mm, slice thickness = 1.0 mm, voxel size = 1.0 × 1.0 × 1.0 mm, and 192 sagittal slices.

### fMRI data preprocessing

fMRI data were preprocessed using the fmriprep pipeline^62^ (version 20.2.5), an open-source tool developed for automated preprocessing of fMRI data. A series of standard preprocessing steps were applied with the default settings and achieved through a combination of ANTs (http://stnava.github.io/ANTs/), FSL^63^, and FreeSurfer (http://surfer.nmr.mgh.harvard.edu/). Specifically, T1-weighted structural images underwent brain extraction, tissue segmentation, and spatial normalization to the MNI152NLin2009cAsym template. Functional images were slice-timing corrected, head motion corrected, susceptibility-derived distortion corrected, coregistered to T1-weighted images, and resampled to the standard space. No spatial smoothing was applied.

### ROI definition

We selected three auditory ROIs: the Heschl’s gyrus (HG), the planum temporale (PT) and the posterior superior temporal gyrus (pSTG). The three ROI were defined by the Harvard-Oxford atlas^64^. We used the probabilistic atlas thresholded at 50%. We selected four visual ROIs: V1, V4, the lateral occiptal cortex (LOC), and the fusiform gyrus (FG). The LOC and FG were also defined by the Harvard-Oxford atlas. Specifically, we used the ”inferior division of LOC” and the ”temporal occipital fusiform cortex” to define the LOC and FG. Because the Harvard-Oxford atlas does not include V1 and V4, we used the Julich brain atlas^65^ to define V1 and V4. We again used the probabilistic atlas thresholded at 50%. We also used the Harvard-Oxford atlas to define the hippocampus and the parahippocampal cortex (PHC). The PHC was named as the posterior division of the parahippocampal gyrus in the atlas. We selected the retrosplenial cortex (RSC) and the precuneus (PCu) using the HCP-MMP atlas^66^. The atlases were resampled to match the resolution of the fMRI, and we visually checked to ensure the selected ROIs were correctly localized. All implementations were performed using Nilearn (https://nilearn.github.io/stable/index.html).

### ROI-based fMRI univariate analysis

To estimate task-evoked univariate responses, we applied the general linear model (GLM) using SPM12 (https://www.fil.ion.ucl.ac.uk/spm/software/spm12/). We first smoothed the fMRI data using a 6 mm full-width at half-maximum (FWHM) Gaussian kernel. For each task, we concatenated fMRI data across all runs and modeled the task-evoked responses using a design matrix that included the onset and duration of each task event. These events were convolved with the canonical hemodynamic response function (HRF) to form the regressors of interest. Additionally, we included a separate regressor to account for task-irrelevant events, such as the oddball detection task in the audiovisual perception task and the beep detection task in the visual memory recall task. We also incorporated six motion parameters as nuisance regressors, along with run-specific regressors to account for inter-run variability. A high-pass filter with a cutoff of 128 seconds was applied to remove low-frequency drifts. Following model estimation, we extracted beta estimates and averaged across voxels within a brain ROI. Next, we assessed whether the task-evoked responses within an ROI were significantly greater than zero using one-sample t-tests.

### ROI-based fMRI time-resolved multivariate pattern analysis

To estimate time-resolved brain responses, we applied GLM with a finite impulse response (FIR) model, which estimated neural response at each time bin (TR = 1 second) without assuming the shape of the hemodynamic response function. To enhance statistical power, we concatenated fMRI data across all runs within each task. Given the rapid event-related design, we employed the square-separate method proposed by Turner et al.^42^. Specifically, for each stimulus, we constructed a design matrix comprising 16 regressors of interest, each corresponding to a time point spanning 16 seconds following stimulus onset. Additionally, 16 separate regressors of non-interest were included to model responses to all other stimuli and task events for 16 time points. We further included six motion parameters as nuisance regressors and one run-specific regressor to account for inter-run variability. A high-pass filter with a cutoff of 128 seconds was applied to remove low-frequency drifts. This GLM procedure was repeated for all 80 stimuli, yielding beta weights for 16 time points at a 1-second resolution for each stimulus. We then converted beta estimates into t-values by contrasting them against the implicitly modeled baseline. All GLM analyses were implemented using SPM12 (https://www.fil.ion.ucl.ac.uk/spm/software/spm12/).

Next, we extracted response patterns from t-maps within a brain ROI at each time point and applied multivariate pattern analysis (MVPA) to assess the categorical decoding over time. Specifically, we employed a linear support vector machine (SVM) classifier with a repeated stratified k-fold cross-validation (5 folds and 10 repeats) procedure to estimate decoding accuracy. All multivariate analyses were implemented using Nilearn (https://nilearn.github.io/stable/index.html).

### Cross-correlation analyses

To quantify the temporal differences between regions, we performed cross-correlation analysis on the decoding accuracy time courses. First, the accuracy values were linearly interpolated to 100 ms resolution to increase temporal precision. For each pair of ROIs, we computed the temporal shift that maximized the cosine similarity between their decoding curves. In each comparison, the averaged decoding accuracy curve of one region was treated as the reference, while the accuracy values of the other region were resampled 10,000 times using bootstrap resampling to estimate the median and 95% confidence interval (CI) of the temporal lag. A positive lag indicated that the compared region (non-reference) was delayed relative to the reference region, whereas a negative lag indicated the opposite. Repeating the analysis with the order of the two regions reversed yielded consistent results, confirming that the findings were not dependent on the choice of reference region.

### Statistical analysis

To evaluate task-evoked univariate responses, we conducted one-sample t-tests on the beta estimates extracted and averaged for each ROI. For each task, we tested whether the mean beta estimate across participants in a given ROI was significantly greater than zero, indicating reliable task-related activation. We applied false discovery rate (FDR) correction to account for multiple comparisons across ROIs, ensuring robust statistical inference.

To determine the time points at which decoding accuracy was significantly above the chance level, we first normalized the accuracy values by the mean of the maximum peak across time points. This normalization aimed to control for differences in signal strength across different ROIs. Next, we conducted one-sided sign permutation tests with 10,000 randomizations to assess whether decoding accuracy at each time point exceeded the empirical null distribution. To correct for multiple comparisons across time points, we applied false discovery rate (FDR) correction at q < 0.05, ensuring robust statistical inference.

All statistical analyses were performed using Python (SciPy, StatsModels, and customized codes).

## Data availability

The publicly available data used for analysis are available in the following repositories: https://osf.io/xv6st/overview.

## Code availability

The publicly available codes for analysis are available in the following repositories: https://osf.io/xv6st/overview.

## Acknowledgements

This study was supported by the Canada First Research Excellence Fund (CFREF) through a BrainsCAN grant to Y.M., a Vector Institute Research Grant to Y.M, and an NSERC Discovery Grant to Y.M..

## Author Contributions

Conceptualization: Y.H. and Y.M.; Methodology: Y.H., J.D and Y.M.; Software: Y.H.; Validation: Y.H.; Formal Analysis: Y.H.; Investigation: Y.H., J.D, and Y.M.; Resources: Y.M.; Data Curation: Y.H.; Writing – Original Draft: Y.H.; Writing – Review and Editing: Y.H, J.D. and Y.M.; Visualization: Y.H.; Supervision: J.D. and Y.M.; Project Administration: Y.M.; Funding Acquisition: Y.M.

## Supplementary Information

### Supplemental Table

**Table S1:**
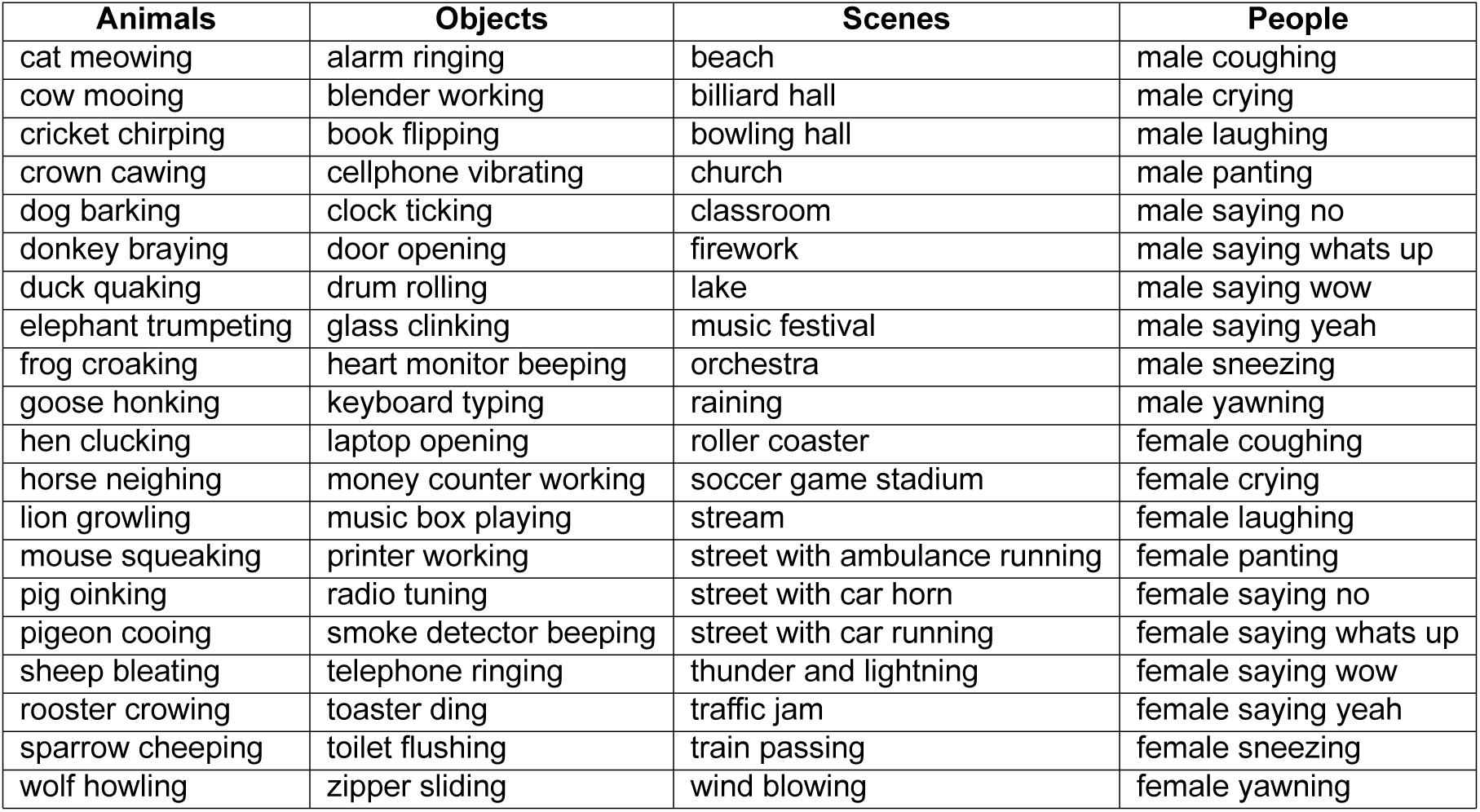
Descriptions of video stimuli spanning categories of animals, objects, scenes, and people. Each line describes the content of one video stimulus. The video contain matching visual and audio for one second.

| **Animals** | **Objects** | **Scenes** | **People** |
| --- | --- | --- | --- |
| cat meowing | alarm ringing | beach | male coughing |
| cow mooing | blender working | billiard hall | male crying |
| cricket chirping | book flipping | bowling hall | male laughing |
| crown cawing | cellphone vibrating | church | male panting |
| dog barking | clock ticking | classroom | male saying no |
| donkey braying | door opening | firework | male saying whats up |
| duck quaking | drum rolling | lake | male saying wow |
| elephant trumpeting | glass clinking | music festival | male saying yeah |
| frog croaking | heart monitor beeping | orchestra | male sneezing |
| goose honking | keyboard typing | raining | male yawning |
| hen clucking | laptop opening | roller coaster | female coughing |
| horse neighing | money counter working | soccer game stadium | female crying |
| lion growling | music box playing | stream | female laughing |
| mouse squeaking | printer working | street with ambulance running | female panting |
| pig oinking | radio tuning | street with car horn | female saying no |
| pigeon cooing | smoke detector beeping | street with car running | female saying whats up |
| sheep bleating | telephone ringing | thunder and lightning | female saying wow |
| rooster crowing | toaster ding | traffic jam | female saying yeah |
| sparrow cheeping | toilet flushing | train passing | female sneezing |
| wolf howling | zipper sliding | wind blowing | female yawning |

